# Targeted genome editing of the non-model cyanobacterium *Cyanothece* PCC 7425 via CRISPR/Cas12a

**DOI:** 10.64898/2026.05.09.723881

**Authors:** Monis Athar Khan, Anne Durand, Fériel Skouri-Panet, Karim Benzerara, Corinne Cassier-Chauvat, Franck Chauvat, Soufian Ouchane

**Affiliations:** Université Paris-Saclay, CEA, CNRS, Institute for Integrative Biology of the Cell (I2BC), Gif-sur-Yvette, France; Sorbonne Université, Muséeum National d’Histoire Naturelle, UMR CNRS 7590. Institut de Minéralogie, de Physique des Matériaux et de Cosmochimie (IMPMC), Paris, France

**Keywords:** non-model cyanobacteria, *Cyanothece*, CRISPR/Cas12a, and gene knockout

## Abstract

Cyanobacteria are diverse photosynthetic microorganisms of great interest for fundamental science and sustainable biotechnological applications. However, their polyploidy makes genetic manipulation challenging and time-consuming. The development of CRISPR/Cas tools has greatly accelerated genome editing and metabolic engineering of few cyanobacterial model species. In this work, we extend the CRISPR/Cas12a system for targeted gene deletion in the non-model cyanobacterium *Cyanothece* PCC 7425, interesting for its ability to perform intracellular calcium carbonate (CaCO_3_) biomineralization, nitrogen fixation, etc. We demonstrate for the first time its tractability to gene knockout by generating deletion mutants of four genes (*cax3-cax4*, *gor*, and *sodB*) acting in metabolism and/or response to stresses, using Cas12a mediated homologous recombination. Importantly, full chromosome segregation was rapidly achieved after a single round of selection in all cases. All mutants were genotypically and phenotypically characterised. Moreover, biochemical analysis in the case of *ΔsodB* mutant further confirmed its targeted deletion. Overall, CRISRPR/Cas12a provides a rapid and efficient system for genome editing in *Cyanothece* PCC 7425, establishing this organism as a versatile model for studying oxidative stress pathways, metal toxicity and moreover, the still poorly known mechanism(s) of intracellular CaCO_3_ biomineralization.

**Key Points:**

- Rapid and efficient CRISPR/Cas12a editing established in *Cyanothece* PCC 7425.
- Fully segregated knockout mutants obtained after single selection round.
- Platform for nuclear waste bioremediation and other biotechnological applications.

## Introduction

Cyanobacteria have been pivotal in shaping Earth’s history and advancing our understanding of fundamental cellular processes. Through the evolution of oxygenic photosynthesis, they contributed majorly in the oxygenation of the atmosphere about ∼2.3 to 2.5 billion years ago during the Great Oxidation Event, dramatically impacting the evolution of life on Earth (Anbar et al., 2007; Dismukes et al., 2001). In comparison to algae and plants, cyanobacteria offer rapid growth and easier genetic manipulation, making them important model organisms for photosynthetic research. The ability of cyanobacteria to utilize water, CO₂, and sunlight as inputs to drive their metabolism positions them as an attractive chassis for “green and sustainable” biotechnological applications like bioremediation and bioproduction. Their remarkable diversity offers a variety of fascinating features (Cassier-Chauvat et al., 2021). Yet, research on cyanobacteria has largely focused on a few genetically tractable model species, including the unicellular cyanobacteria *Synechocystis* PCC 6803, *Synechococcus* PCC 7942, *Synechococcus* PCC 7002, and the filamentous cyanobacterium *Anabaena* PCC 7120 (Elhai and Wolk, 1988; Golden et al., 1987; Kuhlemeier and van Arkel, 1987; Marraccini et al., 1993). Many of the genetic tools developed are restricted to these few species only, while genetic transformation remains difficult for others (Taton et al., 2014). This problem arises from the complex biology of cyanobacteria that poses several difficulties to their genetic editing. In particular, they harbour multiple copies of chromosome (Watanabe, 2020), which makes genetic manipulation time-consuming, as multiple rounds of segregation are needed to ensure genetic modification in all chromosome copies.

The advent of CRISPR/Cas tools has helped overcome these challenges, greatly accelerating the genetic engineering of model cyanobacteria (Behler et al., 2018; Ungerer and Pakrasi, 2016; Wendt et al., 2016). Cas12a (previously Cpf1) from *Francisella novicida* was shown to be far less toxic than Cas9, leading to its development for genetic engineering of cyanobacteria (Ungerer and Pakrasi, 2016; Wendt et al., 2016). Cas12a has additional advantages as compared to Cas9: (i) It creates a staggered cut with 5’ overhang as opposed to a blunt end generated by Cas9, promoting homology directed repair (Chauhan et al., 2023); (ii) Cas12a requires a shorter RNA as compared to dual crRNA-tracrRNA system used by Cas9 (Zetsche et al., 2015); and (iii) Cas12a recognizes a T-rich protospacer-adjacent motif (PAM), expanding its use in AT-rich genomes (Zetsche et al., 2015). The efficiency of the CRISPR/Cas12a system has been demonstrated by engineering marker-less knockouts, knockins and point mutations in *Synechococcus elongatus* UTEX 2973, *Synechocystis* PCC 6803 and *Anabaena* PCC 7120 (Ungerer and Pakrasi, 2016). Since then, CRISPR/Cas12a has been successfully extended to other unicellular and filamentous cyanobacteria (Bandyopadhyay et al., 2024; Gao et al., 2024; Sengupta et al., 2020; Victoria et al., 2024) and is emerging as a widely useful tool for engineering cyanobacteria.

In this study, we extended the application of CRISPR/Cas12a-based gene deletion to the non-model unicellular cyanobacterium, *Cyanothece* PCC 7425 (hereafter, *Cyanothece* 7425). *Cyanothece* 7425 is able to fix nitrogen and generate H_2_ (Bandyopadhyay et al., 2011) and can produce cyanobactins, a class of ribosomal cyclic peptides that are of significant pharmacological relevance (Houssen et al., 2012). It also harbours panoply of antioxidant enzymes like glutathione disulfide reductase (Gor) and two superoxide dismutase (SOD) to deal with oxidative stress. Another salient feature of *Cyanothece* 7425 is its capability to form intracellular amorphous calcium carbonate (iACC) inclusions enclosed in a microcompartment (Benzerara et al., 2022, 2014). This process contributes to active sequestration of alkaline earth elements like calcium, strontium, barium and radium, thus, playing an important role in their biogeochemical cycling and implications for nuclear waste bioremediation (Blondeau et al., 2018; Mehta et al., 2019; Pamart et al., 2025). Several other cyanobacteria from diverse taxa have also been shown to form iACC inclusions (Benzerara et al., 2022, 2014). Moreover, the precipitation of calcium carbonate (CaCO_3_) by cyanobacteria also offers the potential for carbon capture and sequestration to reduce CO_2_ emissions (Jansson and Northen, 2010). However, the lack of genetic tools in iACC-forming cyanobacteria has limited progress in elucidating the mechanisms underlying iACC formation. Building on previous reports of RSF1010-derived plasmid transfer for gene overexpression (Chenebault et al., 2020), in this work we used another RSF1010-derived plasmid, pSL2680 (Ungerer and Pakrasi, 2016), to develop targeted gene deletion in *Cyanothece* 7425 via CRISPR/Cas12a. We successfully deleted genes (*cax3-4*, *gor* and *sodB*) of varying length, including a ∼3.7 kb deletion spanning two adjacent genes.

## MATERIALS AND METHODS

### Strains and culture conditions

All strains of *Cyanothece* 7425 were cultured at 30°C under continuous agitation at 140 rpm (Infors rotary shaker) and standard light condition: cool white light (Sylvania Luxline Plus) at intensity of ∼2500 lux (32 µmol m^−2^s^−1^). Cells were grown in Mineral Medium (MM) (Domain et al., 2004), a modified version of the standard BG-11 medium (Stanier et al., 1971), containing 1.89 mM of sodium carbonate (Na₂CO₃) as the carbon source. For growth on plate, MM was solidified with 1% Bacto Agar (Difco). *E. coli* strain NEB 10-beta (New England Biolabs) was used for cloning and propagation of plasmids. The helper strain *E. coli* CM404 (Marraccini et al., 1993) was used for triparental conjugation. *E. coli* strains were grown in liquid or solid LB medium at 30°C (CM404) or 37°C (NEB 10-beta). The concentration of antibiotics used for selection were ampicillin (Amp) 100 µg/mL, kanamycin (Km) 50 µg/mL, streptomycin (Sm) 25 µg/mL and spectinomycin (Sp) 75 µg/mL for *E. coli* and (Sm) 5 µg/mL, (Sp) 5 µg/mL and (Km) 25 µg/mL for *Cyanothece* 7425.

### Plasmid construction

All cloning steps were carried out in the *E. coli* strain NEB10-beta. The primers are listed in Table S2. All enzymes used were from New England Biolabs (NEB) except AarI (Thermo Fisher Scientific). The plasmids were sequenced using the Mix2Seq kit (Eurofins Genomics). All plasmids used are listed in Table S1.

### pSL2680-trCas12a vector construction

The vector pSL2680 (Addgene plasmid # 85581) was a gift from Himadri Pakrasi (Ungerer and Pakrasi, 2016). The vector was digested with EcoRI and BspEI restriction enzymes followed by blunting using DNA Polymerase I, Large (Klenow) Fragment and ligation using T4 DNA ligase. The consequent vector pSL2680-Δcas12a, had ∼1.8 kb fragment of *cas12a* gene removed resulting in truncated Cas12a (Fig. S1).

### Editing vector construction

For the construction of vector pSL-*Δcax3-4*, first the complementary phosphorylated oligos cr-cax3-4_Fw/Rv (Integrated DNA Technologies) corresponding to the designed crRNA were annealed by heating at 65 °C. The annealed oligos were ligated at the AarI-linearized pSL2680 vector, generating vector pSL-crRNA-cax3-4. After successful cloning of the spacer, the homology platforms of ∼600 bp were generated by overlapping PCR (Phusion polymerase, NEB) on genomic DNA of *Cyanothece* 7425 using oligos F1-cax_Fw/Rv and F3-cax_Fw/Rv. The Sm^R^Sp^R^ cassette was obtained by PCR using oligos F2-cax_Fw/Rv on purified pC plasmid (Chenebault et al., 2020). The three fragments were cloned at the SalI site in pSL-crRNA-cax3-4 by HiFi DNA assembly (NEB), generating the final vector pSL-*Δcax3-4* (Fig. S3).

The vector pSL-*Δgor* was constructed in a similar way except the homology repair template (∼500 bp homology platforms) was recovered from synthesized plasmid pT-*Δgor* (Twist Bioscience, Table S1) using oligos F-pT-Δgor_Fw/Rv (Fig. S5).

The vector pSL-*ΔsodB* (Fig. S6) was constructed as described for pSL-*Δcax3-4*, using the respective primers listed in Table S2.

### Transfer of plasmids via triparental conjugation

The plasmids were transferred from *E. coli* to *Cyanothece* 7425 via triparental conjugation as described previously (Chenebault et al., 2020). Five mL cultures of *E. coli* CM404 and NEB 10-beta (harbouring the desired plasmid) were grown overnight as described above. Next day, the cultures were washed with LB twice and then resuspended to 1.3 x 10^9^ cells/mL final concentration (*E. coli* OD_600nm_ 1 = 2 x 10^9^ cells/mL). *Cyanothece* 7425 was grown as described above until OD_750nm_ ∼0.6- 1 and washed twice with MM. Then cells were resuspended to a final concentration of 1.25 x 10^7^ cells/mL (*Cyanothece 7425* OD_750nm_ 1 = 2.5 x 10^7^ cells/mL). Then, 100 µL of this suspension was mixed with 30 µL suspension each of *E. coli* CM404 and *E. coli* NEB 10-beta carrying the desired plasmid (final ratio of *E. coli*: *Cyanothece 7425* = 3:1). For negative control, 100 µL of *Cyanothece 7425* were mixed with 60 µL LB medium to monitor the frequency of spontaneous antibiotic-resistant mutants. Finally, 30 µL aliquots of these mixtures were spotted on non-selective MM agar plates and incubated at 30°C for 72 hours under standard light (2500 lux). Following incubation, each spot was collected in 100 µL MM and spread onto selective MM plates containing either SmSp 5 µg/mL or Km 25 µg/mL and incubated for 2-3 weeks under standard light conditions.

### Construction of mutant strains

The editing vectors were transferred to *Cyanothece* 7425 via triparental conjugation as described above. After 2-3 weeks of growth, all antibiotic-resistant clones were patched onto selective medium containing 5 µg/mL of both Sm and Sp. Their genomes were analysed by PCR to confirm deletion of the studied genes and full segregation of their chromosome. At least two clones were then grown in liquid MM medium containing SmSp 5 µg/mL for another round of PCR verification and sequencing (Eurofins Genomics).

### SEM/EDXS analysis of wild-type and *Δcax3-4*

Scanning electron microscopy (SEM) was performed at the IMPMC microscopy platform (Sorbonne Université, France), as described previously (Benzerara et al., 2014). Briefly, the wild-type and *Δcax3-4* strains were cultured in the BG11 medium until exponential phase (OD_750nm_ ∼ 0.9 to 1.2). Then 0.5 mL of cell suspension was collected onto 0.22 µm-pore-size polycarbonate filters (Millipore) and rinsed three times with ultrapure water. The filters were air-dried at room temperature and mounted on aluminium stubs using double-sided carbon tape. A thin film of carbon was deposited on the filters using a Leica EM SCD500 evaporator to enhance surface electron conductivity. The SEM apparatus was a Zeiss™ Ultra 55 microscope equipped with a field emission gun. Images were captured in the backscattered electron (BSE) detection mode using an angle-selective backscatter (AsB) detector. The acquisition conditions included an electron accelerating voltage set to 15 kV, a working distance of 7.5 mm, and a 60 μm aperture. Chemical mapping was conducted using energy dispersive X-ray spectrometry (EDXS). Data was processed with the ESPRIT software (Bruker). SEM-EDXS analysis was replicated twice with independent cultures and at least three different areas observed each time.

### Spot viability assay

For spot viability assay, the wild-type and mutant strains were grown in liquid MM until exponential phase (OD_750nm_ ∼0.8 - 1.2). The cultures were centrifuged at 4500 *g* for 10 minutes and washed twice with sterile ultrapure water. The dilution series began with an OD₇₅₀ of 0.1, followed by five rounds of five-fold dilution. A drop of 10 µL from each dilution was spotted onto either MM agar plates or MM agar plates supplemented with different final concentrations (250 µM, 2.5 mM and 25 mM) of CaCl_2_ for *Δcax3-4* and CuCl_2_ (5 µM), CdCl_2_ (2.5 µM) and CoCl_2_ (5 µM) for *Δgor*. The plates were incubated for 14 days under standard light at 30°C and imaged after.

### SOD *in-gel* activity assay on non-denaturing gel

50 mL cultures of wild type (WT) and *ΔsodB* strains were grown until late exponential phase (OD_750nm_ = ∼1.5-1.8). Cells were prepared following the previously described protocol (Ivleva and Golden, 2007). Protein concentration was determined by BCA using BSA as standard. Then 30 µg of total protein extracts was separated on a 16% non-denaturing polyacrylamide gel and stained for SOD activity as described (Weydert and Cullen, 2010). Gel was first incubated in 20 mL solution of 50 mM Tris-HCl (pH = 7.2) with 0.85% TEMED (Sigma-Aldrich) and 56 µM Riboflavin-5-Phosphate (Sigma-Aldrich) for 15 min in light at room temperature (RT), followed by the addition of 2 mg/mL Nitroblue Tetrazolium Chloride (Sigma-Aldrich) and 15 min incubation in the dark at RT. Gel was washed twice in ultrapure water and left at RT in a light box until SOD-positive staining appeared.

### Disc diffusion assay

500 µL (OD_750nm_ = 0.8) of WT or *ΔsodB* were mixed with 6 mL semi-solid MM-agar (50%) and uniformly spread on MM-solidified agar plates. 6 mm sterile discs were soaked with 5 μL of 5 mM menadione (Sigma-Aldrich) and placed in the middle of the plate. Plates were incubated at 30°C under standard light condition for 7 days prior to photography.

### Verification of absence of editing plasmid

To check if the mutant strains have lost the editing vector, the WT, mutant strains and Km^R^ and Sm^R^Sp^R^ positive control strains were grown in liquid medium until reaching mid-exponential phase. Then, 10 µL of each culture was patched on MM agar plates containing either SmSp 5 µg/mL or Km 25 µg/mL. The plates were then incubated for 10 days under standard light at 30°C and imaged after. Cells collected from the plates were then used for PCR verification with primers targeting the editing vector (See Table S2).

## RESULTS

### Toxicity of Cas12a in *Cyanothece* PCC 7425

First, the potential toxicity of Cas12a expression in *Cyanothece* 7425 was assessed by introducing into it the pSL2680 vector (Ungerer and Pakrasi, 2016) via triparental conjugation as we previously described (Chenebault et al., 2020). On several attempts, very few Km^R^ clones (2 to 6 clones) were obtained with pSL2680 as compared to the large number of clones (>200) obtained with another RSF1010-derived replicative plasmid pSB2T (Marraccini et al., 1993) used as positive control (Fig. 1, top panel). The low number of clones obtained with pSL2680 suggested that the expression of Cas12a might be toxic to *Cyanothece* 7425. To test this hypothesis, a ∼1.8 kb segment from the ∼3.9 kb coding sequence of the *cas12a* gene was removed from the pSL2680 vector, yielding the pSL2680-trCas12a plasmid encoding a truncated Cas12a protein. The trCas12a variant missing the Cas12a amino acids 733-1330, lacks the RuvC and nuclease domain and hence cannot mediate DNA cleavage (Swarts, 2019). Both pSL2680-trCas12a (Fig. S1) and pSL2680, were then conjugated into *Cyanothece* 7425. As expected, conjugation of pSL2680-trCas12a vector produced much more clones (>100), comparable to the positive control vector pSB2T (Fig. 1, bottom panel), confirming the toxicity of Cas12a. Nevertheless, despite its toxicity, the recovery of clones harbouring the pSL2680 plasmid (Fig. 1, top panel) indicated that the CRISPR/Cas12a system could be used to generate deletion mutants.

**Figure 1:**
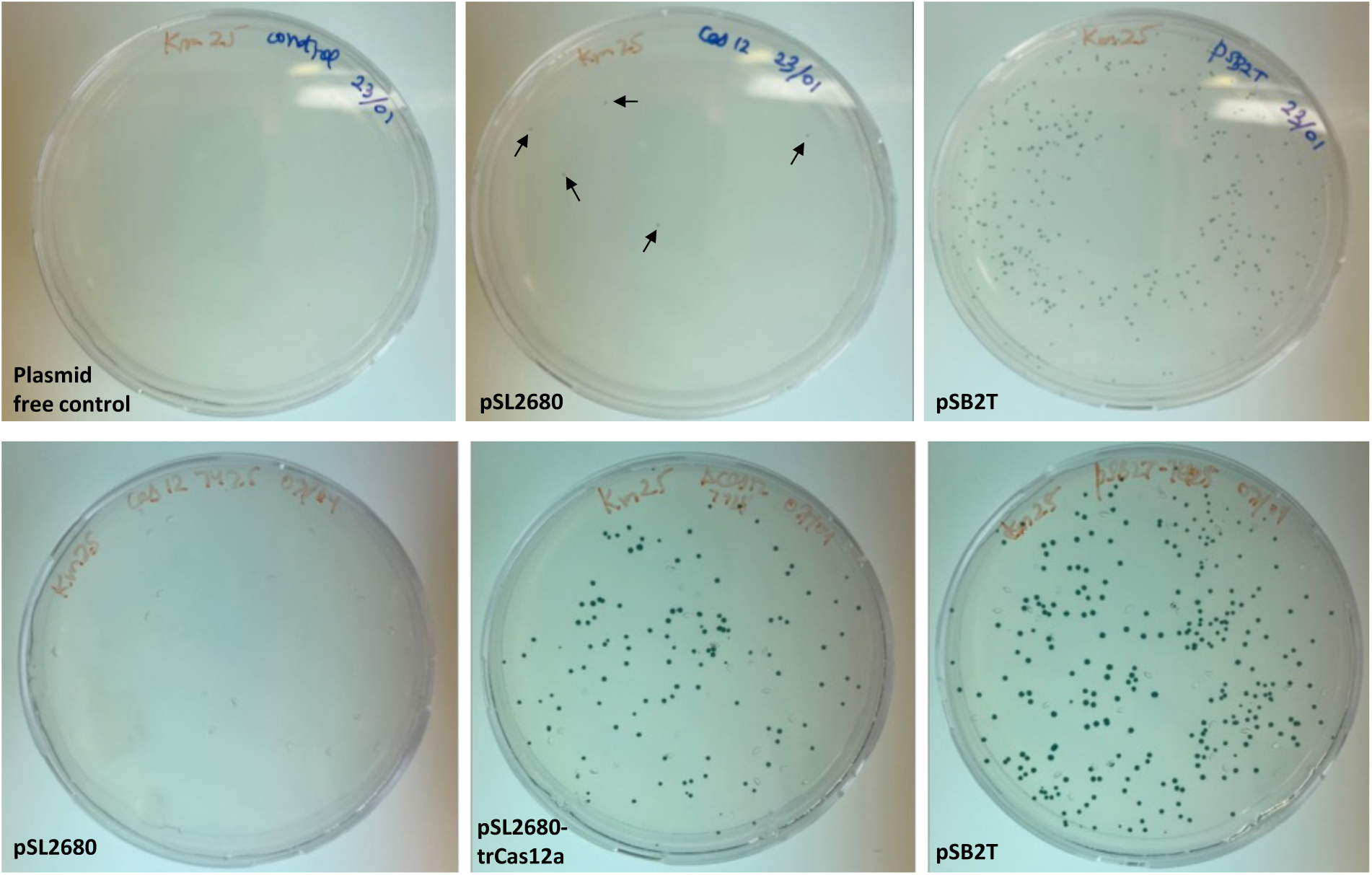
Conjugative transfer of plasmids into *Cyanothece* PCC 7425. Photographs of kanamycin containing plates resulting from the conjugation with the plasmids pSB2T (positive control), pSL2680 or its pSL2680-trCas12 derivative constructed in this work, or no plasmid (negative control for checking possible Km^R^ spontaneous mutants). The few Km^R^ clones obtained with pSL2680 vector (attesting the toxicity of Cas12) are indicated with arrows. The plates were imaged after 2 (upper row) or 3 (lower row) weeks of incubation.

### CRISPR/Cas12a mediated deletion of antioxidant enzymes glutathione reductase and superoxide dismutase in *Cyanothece* PCC 7425: Proof of concept

Cyanobacteria are frequently exposed to oxidative stress when performing oxygenic photosynthesis, which together with cellular respiration is a major source of reactive oxygen species (ROS) (Latifi et al., 2009). Under high light stress, increased amount of ROS is produced leading to inactivation of the photosynthetic machinery (Nishiyama et al., 2001). Among ROS, the superoxide anion (O₂•⁻) is detoxified through the action of superoxide dismutase (SOD) that transforms it into H_2_O_2_ and O_2_ and the H_2_O_2_ produced is further converted to H_2_O by catalases and peroxidases. Among the latter enzymes, glutathione peroxidases (Gpxs) reduce H_2_O_2_ to H_2_O using electrons provided by the antioxidant tripeptide glutathione (GSH) as a cofactor (Ursini et al., 1995). In this process, GSH is oxidized to glutathione disulfide (GSSG), which can be reduced to GSH by the NADPH-dependent enzyme glutathione reductase (Gor). Thus, Gor plays an important function in maintaining the cellular GSH pool. Thus, both Gor and SodB enzymes are critical for defence against oxidative stress in many organisms including cyanobacteria (Cameron and Pakrasi, 2010; Cassier-Chauvat et al., 2021; Latifi et al., 2009). To validate the potential use of CRISPR/Cas12a as a genetic tool to generate mutants in *Cyanothece* 7425, we used it to delete the *gor* and *sodB* genes.

#### Deletion of the gor gene

Previous attempts to delete the *gor* gene (KEGG ID: Cyan7425_3289) in *Cyanothece* 7425 using classical (CRISPR-independent) gene replacement via natural transformation failed (unpublished data). Consequently, in this study we used the CRISPR/Cas12a system to delete the single *gor* gene in *Cyanothece* 7425. An important determinant of successful gene deletion using the CRISPR/Cas12a system, is the sequence of PAM site of the CRISPR target (Niu et al., 2019). These authors found that CRISPR targets with the PAM site 5’-TTTV-3’ (where V= A, G or C) exhibited a higher editing efficiency (95.2%) as compared to those with a 5′-GTTV-3′ PAM (75.9%). Therefore, for the deletion of *gor* gene, we selected the PAM site 5’-TTTG-3’ (Fig. 2A). For the recombination repair template, homology arms of ∼500 bp upstream of the *gor* start codon and downstream of its stop codon, flanking a spectinomycin/streptomycin resistance (Sm^R^Sp^R^) marker were chosen (Fig. 2B). Hence, if the CRISPR/Cas12a system functions as intended, it would enable the full replacement of *gor* gene with the Sm^R^Sp^R^ cassette (Fig. 2B). The crRNA and the repair template were cloned into the pSL2680 vector to generate pSL-*Δgor*, which was subsequently introduced into *Cyanothece* 7425 via triparental conjugation. Selection was carried out on MM medium containing 5 µg/mL of both Sm and Sp. Upon conjugation of the editing plasmid pSL-*Δgor* (Fig. S5), we obtained six clones that were patched onto selective medium containing 5 µg/mL of both Sm and Sp (Fig. 2C). PCR verification showed the clear presence of the mutant chromosome copies in all the clones and the correct integration of the Sm^R^Sp^R^ cassette in place of the target *gor* gene (Fig. 2D, PCR- A and B). However, PCR analysis with primers specific for the *gor* gene revealed that only three (clone n° 2, 4 and 6) of the six clones showed complete absence of WT chromosome copies (Fig. 2D, PCR-C). The presence of WT chromosome copies in other clones (clone n° 1, 3 and 5) suggests that chromosome segregation has not yet been completed. For further phenotypic analysis one fully segregated clone (clone n° 2) was chosen.

**Figure 2:**
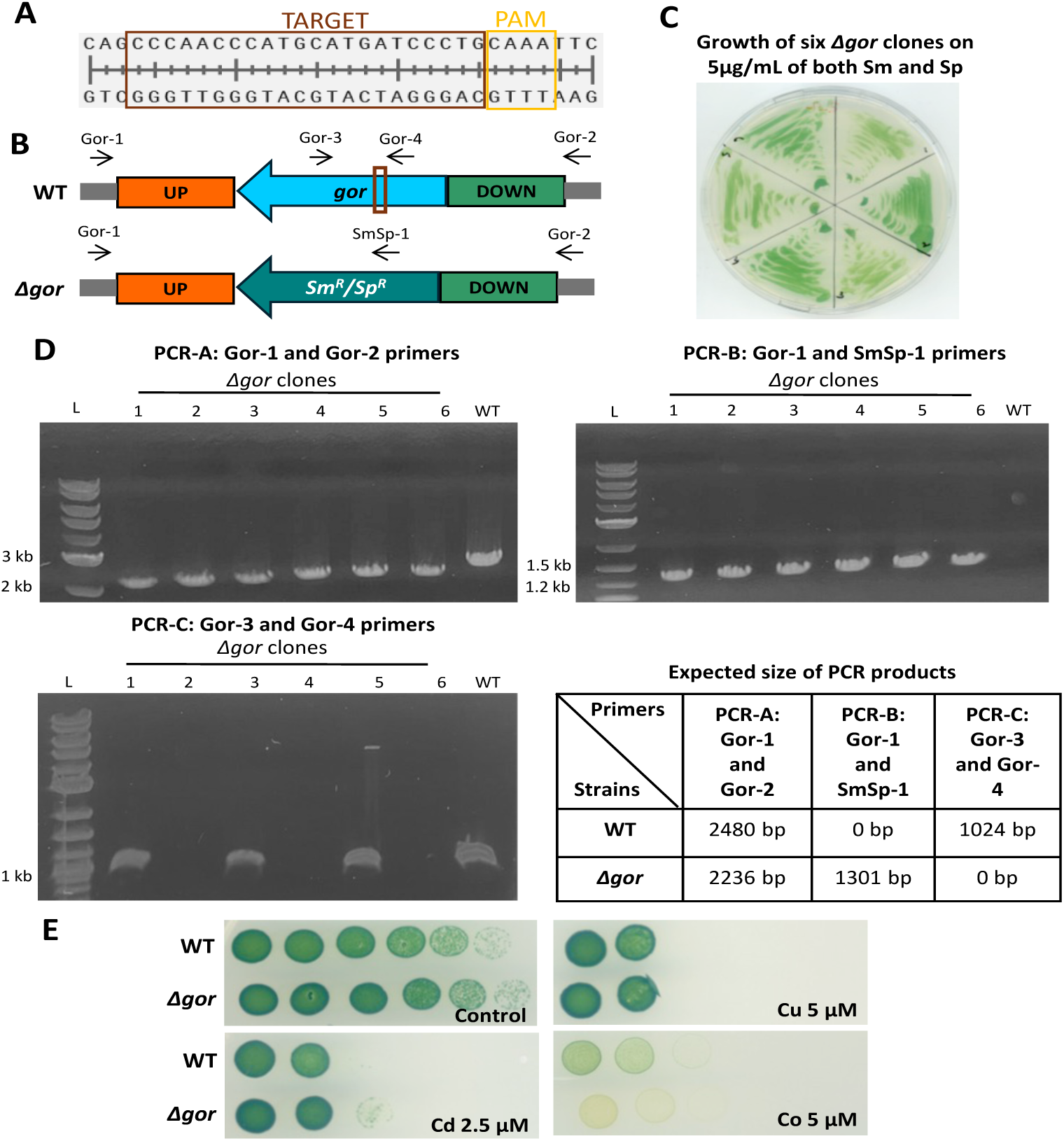
Targeted deletion of the *gor* gene in *Cyanothece* PCC 7425. **(A)** Nucleotide sequence of the crRNA showing the PAM site and the Cas12a target region. **(B)** Schematic representation the wild-type (WT) and mutant (*Δgor*) chromosome. The genes are represented by large arrows. The small black arrows represent the primers used for verification (Table S2) and the brown box indicates the target site of the crRNA. **(C)** Representative image of six *Δgor* clones grown for 10 days on plates containing Sm and Sp. **(D)** Three PCR strategies were employed to confirm successful deletion of the *gor* gene in six Sm^R^Sp^R^ clones, showing complete absence of WT chromosome copy in the *Δgor* clone n° 2,4 and 6. In *Δgor* clone n° 1, 3 and 5, the WT chromosome copy was detected suggesting segregation has not yet been completed. PCR products (expected sizes indicated in table) were resolved on 1% agarose gels. L denotes DNA ladder (1 kb plus, NEB). **(E**) For spot viability assay, cell cultures of WT and *Δgor* strain were adjusted to OD₇₅₀ = 0.1, followed by 1:5 serial dilution prior to spotting as 10 µL drops onto either solid MM medium (control) or containing the indicated metal concentration in excess. The plates were incubated for 14 days at 30°C under standard light and imaged after.

Under standard photoautotrophic conditions, the WT and *Δgor* mutant showed no growth differences (Fig. 2E). In cyanobacteria, the glutathione system has been shown to contribute to tolerance against metal stress (Cassier-Chauvat et al., 2023). Therefore, we compared the growth of WT and the *Δgor* mutant in presence of toxic concentrations of cadmium (Cd), copper (Cu), and cobalt (Co) with a spot viability assay. As observed for the WT strain, the growth of *Δgor* mutant was not impaired by both Cd and Cu (Fig. 2E). In contrast, while Co moderately affected the growth of the wild type, it induced severe growth defect in the *Δgor* mutant (Fig. 2E).

#### Deletion of the sodB gene

Turning to the SOD family of anti-oxidant enzymes, it is known that depending on the metal co-factor present at their active site, SODs can be categorized into four different isoforms: MnSODs containing manganese, FeSODs containing iron, NiSODs containing nickel and CuZnSODs containing copper/zinc (Boden et al., 2021). The different SODs also vary in terms of their subcellular localization. The genome of *Cyanothece* 7425 encodes two superoxide dismutases, a putative membrane-bound MnSOD (*sodA*; KEGG ID: Cyan7425_3014) and a cytosolic FeSOD (*sodB*; KEGG ID: Cyan7425_1660) (Fig. S2). Given the central housekeeping role of *sodB* in the cytoplasm, where it is primarily responsible for scavenging superoxide radicals generated mainly at photosystem I and protecting cells from damage to 4Fe-4S cluster containing proteins (Herbert et al., 1992; Keyer and Imlay, 1996), we targeted the *sodB* gene in *Cyanothece* 7425 to further establish CRISPR/Cas12a system for gene deletion in this species. The CRISPR target and the editing plasmid were designed as described above for the *gor* gene, with the repair template containing homology arms of ∼500 bp upstream of the *sodB* start codon and downstream of its stop codon, flanking a Sm^R^Sp^R^ marker (Fig. 3A). Following conjugation of the plasmid pSL-*ΔsodB*, we obtained only a single clone which was found to be fully segregated and the Sm^R^Sp^R^ cassette integrated in place of the *sodB* gene (Fig. 3B, PCR-A and B). Furthermore, PCR with primers specific for the *sodB* gene resulted in no amplification in the *ΔsodB* mutant (Fig. 3B, PCR-C). This suggests that SodB is dispensable for the growth of *Cyanothece* 7425.

**Figure 3:**
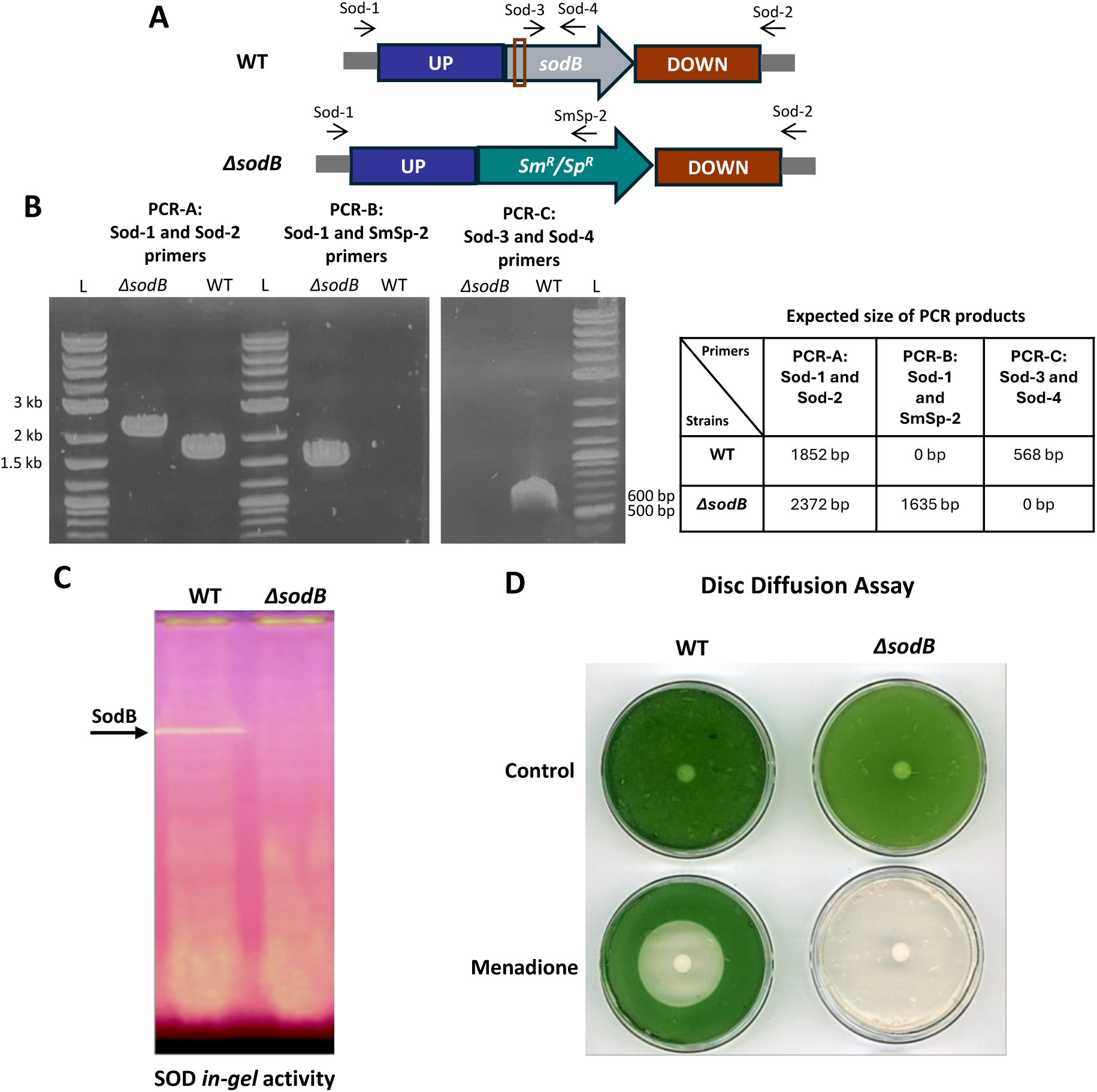
Targeted deletion of the *sodB* gene in *Cyanothece* PCC 7425. **(A)** Schematic representation the wild-type (WT) and mutant (*ΔsodB*) chromosome. The genes are represented by large arrows pointing in direction of their transcription. The small black arrows represent the primers used for verification (Table S2) and the brown box indicates the target site of the crRNA. **(B)** Three PCR strategies were employed to confirm successful deletion of the *sodB* gene, showing complete absence of WT chromosome copy in the *ΔsodB* mutant. PCR products (expected sizes indicated in table) were resolved on 1% agarose gels. L denotes DNA ladder (1 kb plus, NEB). **(C)** The SOD *in-gel* activity of WT and *ΔsodB* was performed with 30 µg total protein extracts loaded on 16% native polyacrylamide gel. The band corresponding to SodB activity is indicated by the arrow. **(D)** Disc diffusion assay of 5 mM menadione sensitivity in the WT and *ΔsodB* strains. The plates were incubated for 7 days at 30°C under standard light and imaged after.

A practical advantage of targeting the *sod* genes is that the SOD enzymatic activity can be readily detected using an *in-gel* colorimetric assay. *In-gel* SOD activity assay revealed a single band in the wild-type strain, which was completely absent in the *ΔsodB* mutant (Fig. 3C). Moreover, we assessed the susceptibility of the *ΔsodB* mutant to menadione using disc diffusion assay. For the wild-type strain, a growth inhibition zone of 15.1 mm in diameter was observed in presence of 5 mM menadione. In contrast, the growth of *ΔsodB* mutant was completely abolished, indicating its hypersensitivity to menadione (Fig. 3D).

### Deletion of two Ca^2+^/H^+^ antiporter encoding *cax* genes: Are *cax* genes involved in iACC formation in *Cyanothece* PCC 7425?

Although the genetic determinants of iACC formation remains unknown, some proteins related to calcium transport have been identified (Bruley et al., 2025). Among calcium transporters, the Ca²⁺/H⁺ antiporter (Cax) appears to be an interesting candidate. In cyanobacteria, Cax proteins are usually involved in exporting Ca^2+^ from the cytoplasm to the periplasm, playing an important role in Ca^2+^ homeostasis (Waditee et al., 2004). Interestingly, the expression of one of the *cax* genes was directly related to biomineralization of CaCO_3_ in the coccolithophore microalga *Emiliania huxleyi*, where it is suggested to transport Ca^2+^ into the intracellular biomineralization compartment known as the coccolith vesicle (Mackinder et al., 2011). Therefore, we hypothesized that the *cax* genes in *Cyanothece* 7425 might be involved in Ca^2+^ trafficking within the bacterium contributing to iACC formation.

The genome of *Cyanothece* PCC 7425 contains four *cax* genes (KEGG ID: Cyan7425_0927, Cyan7425_1779, Cyan7425_3000 and Cyan7425_3001, named *cax1-4*, respectively). Amongst them, *cax3* (Cyan7425_3000) and *cax4* (Cyan7425_3001) are located in a genomic region that also harbours other genes potentially involved in Ca^2+^ transport (Fig. S2)., the adjacent *cax3* and *cax4* genes were chosen as primary targets, as their deletion was expected to perturb Ca^2+^ homeostasis and hence may affect iACC formation. For the recombination repair template, homology arms of ∼600 bp lying both upstream the start codon of *cax3* and downstream the stop codon of *cax4* genes, flanking a spectinomycin/streptomycin resistance (Sm^R^Sp^R^) marker were chosen (Fig. 4A). The editing plasmid pSL-*Δcax3-4* was subsequently introduced into *Cyanothece* 7425 via triparental conjugation. After three weeks, eight colonies were obtained and subsequently patched onto selective medium. All colonies grew well under standard conditions (Fig. 4B) and were subjected to PCR verification to check for successful deletion of *cax3* and *cax4* genes. As shown in Figure 4C, all tested clones harboured only mutant chromosome copies, with no WT chromosome detected confirming *cax3-4* deletion (PCR-A) upon their replacement by the Sm^R^Sp^R^ cassette (PCR-B). To further confirm complete segregation of the chromosome, PCR with primers targeting the *cax4* gene confirmed that in seven out of eight clones, the *cax3-4* genes were fully deleted (Fig. 4C, PCR-C). Overall, these results indicate that the *cax3* and *cax4* genes are not vital for growth under standard conditions.

**Figure 4:**
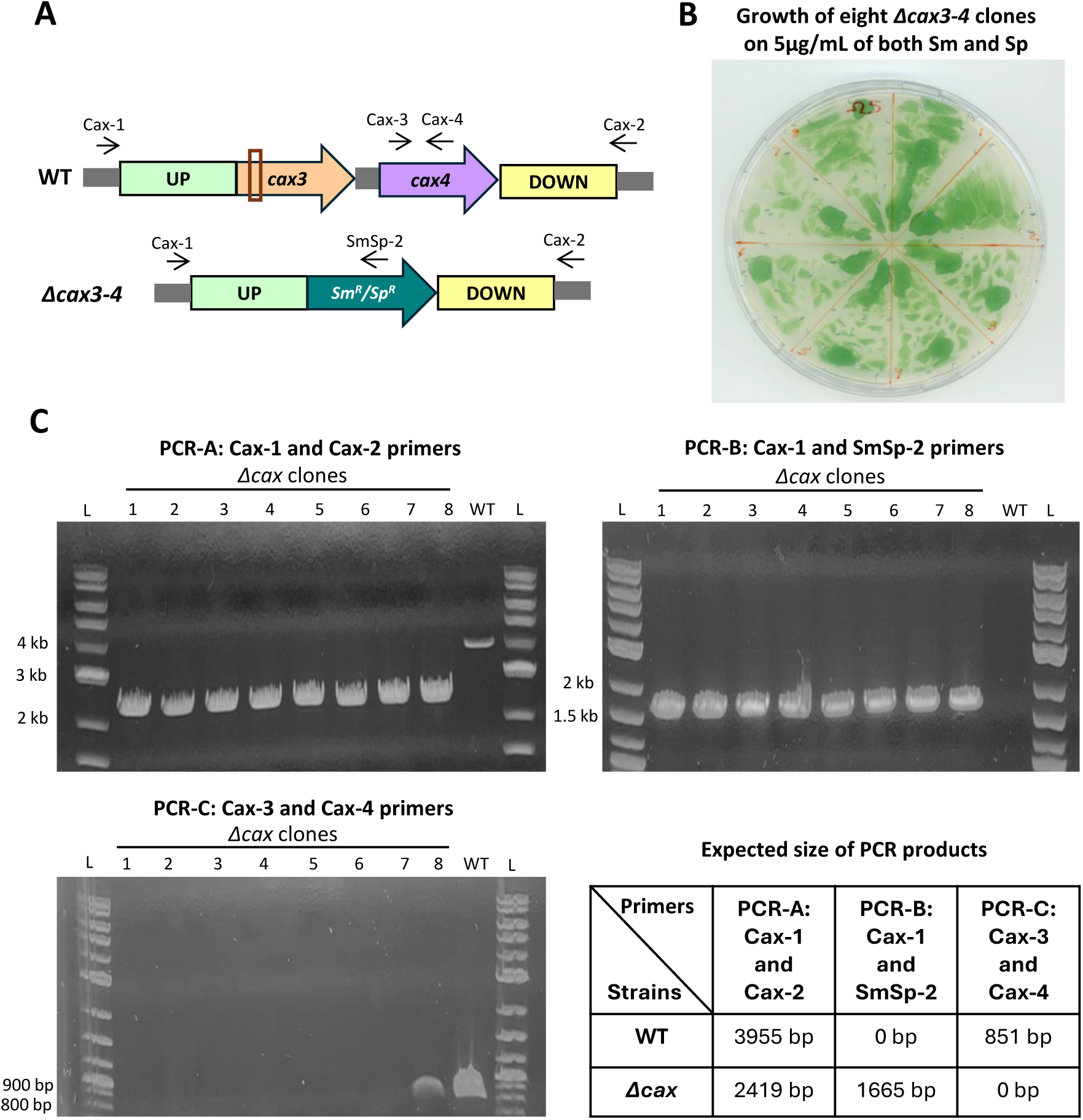
Targeted deletion of *cax3* and *cax4* genes in *Cyanothece* PCC 7425. **(A)** Schematic representation the wild-type (WT) and mutant (*Δcax3-4*) chromosome. The genes are represented by large arrows pointing in direction of their transcription. The small black arrows represent the primers used for verification (Table S2) and the brown box indicates the target site of the crRNA. **(B)** Representative image of eight *Δcax3-4* clones grown for 10 days in presence of both Sm and Sp. The plates were imaged 10 days following incubation. **(C)** Three PCR strategies were employed to confirm successful deletion of the *cax3* and *cax4* genes in eight Sm^R^Sp^R^ clones, showing complete absence of WT chromosome copy in the *Δcax3-4* clone n° 1-7. In *Δcax3-4* clone n° 8, WT chromosome copy was detected suggesting segregation has not yet been completed. PCR products (expected sizes indicated in table) were resolved on 1% agarose gels. L denotes DNA ladder (1 kb plus, NEB).

To assess the impact of *cax3* and *cax4* knockout on iACC formation, one fully segregated clone (clone n°1) was selected and SEM-EDXS analysis was conducted. No significant difference in iACC inclusions was observed between multiple cells of the WT and the *Δcax3-4* mutant (Fig. S4A), suggesting that these *cax3-4* genes are not needed for iACC formation. Next, to assess whether *cax3-4* might be involved in Ca^2+^ export from the cell, we performed a spot viability assay to compare the growth of WT and the *Δcax3-4* mutant under high calcium conditions. Even at 25 mM Ca^2+^ (about 100 times the normal concentration in standard MM medium) no significant difference in growth was observed between the WT and mutant strains (Fig. S4B).

### Verification of absence of the pSL2680-derived editing plasmids in the deletion mutants

To determine whether the deletion mutants have been cured of their respective pSL2680-derived editing plasmid, these strains were patched onto selective media containing either both streptomycin and spectinomycin or kanamycin. Since the backbone of the editing pSL2680 plasmid carries a kanamycin resistance marker (Fig. S1), the loss of the plasmid results in an inability of the mutant strains to grow in presence of kanamycin. Consistently, all three mutants (*Δcax3-4, Δgor* and *ΔsodB*) failed to grow in the presence of kanamycin (Fig. 5). In addition, PCR verification using primers specific for the editing plasmids was negative, confirming the loss of these editing plasmids (Fig. 5).

**Figure 5:**
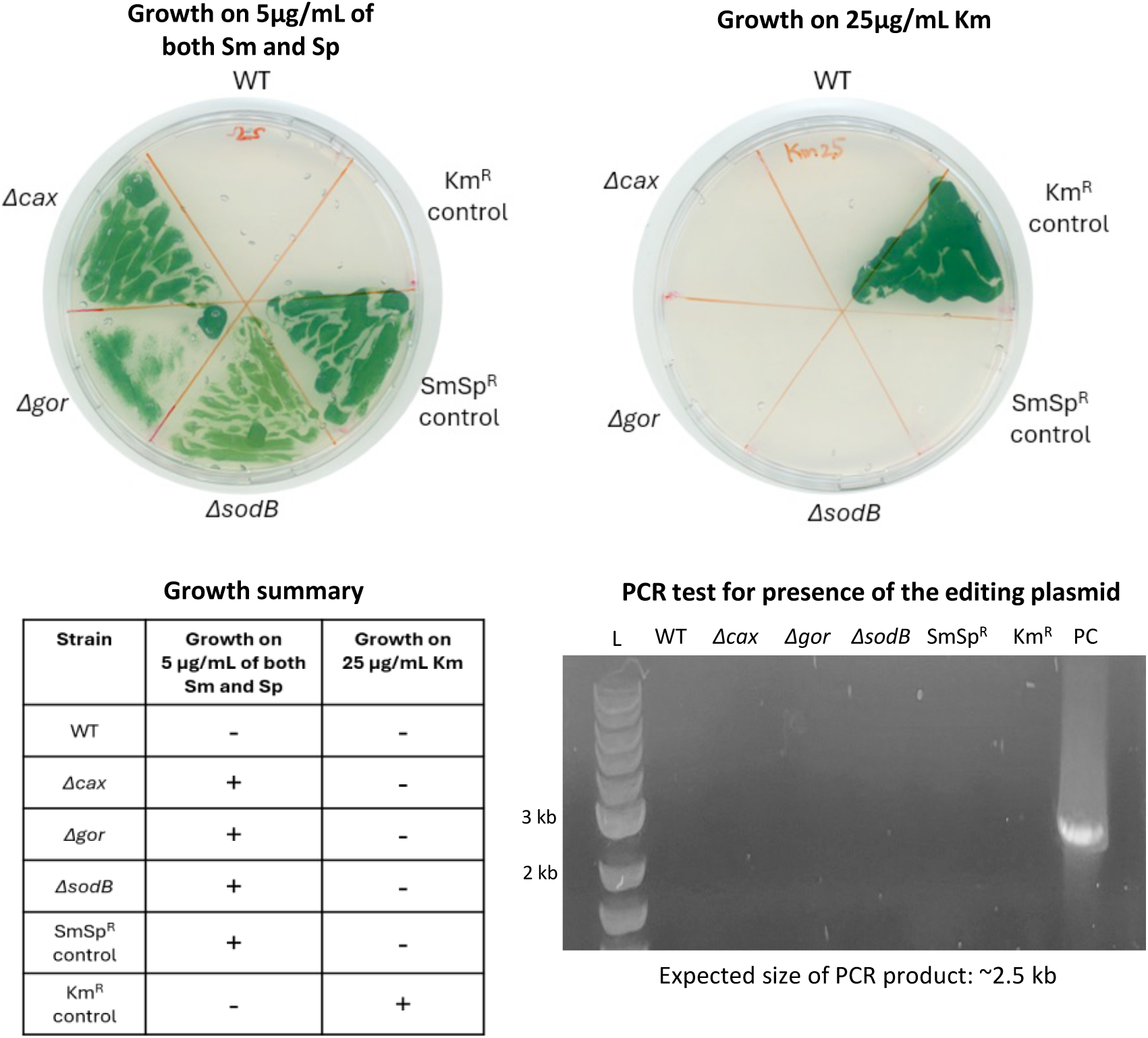
Verification of presence of the editing plasmid. The growth of the indicated strains was assessed on selective media containing either streptomycin and spectinomycin (left plate) or kanamycin (right plate). The table summarises whether each strain exhibited growth (+) or no growth (-) on the respective selective media. PCR analysis using primers (Table S2) targeting the editing vector showing the products resolved on 1% agarose gels. PC is positive control. L denotes DNA ladder (1 kb plus, NEB).

## DISCUSSION

The inability to obtain a high number of colonies with the vector pSL2680 suggested that Cas12a is toxic and causes cell death of *Cyanothece* 7425. It has been proposed that Cas9 toxicity in cyanobacteria could arise from the off-target cleavage of DNA, that cannot be efficiently repaired, ultimately causing cell death (Wendt et al., 2016). The toxicity of Cas12a observed in *Cyanothece* 7425 can be attributed to similar factors. Our finding is consistent with few other studies that reported the toxicity associated with high expression of dCas12a (deactivated Cas12a, that binds to DNA hindering gene expression) (Knoot et al., 2020; Xie et al., 2025). Nevertheless, Cas12a-associated toxicity could be mitigated by employing inducible promoters to fine-tune its expression, which may improve overall gene-editing efficiency (Victoria et al., 2024). Despite the observed toxicity, CRISPR/Cas12a system is rapid and efficient for genome editing in *Cyanothece* 7425. The targeted genes can be replaced with Sm^R^Sp^R^ cassette through homologous recombination following the double-stranded break generated by Cas12a at the desired genomic locus. We show that homology arms of approximately ∼500 bp are sufficient for homologous recombination, which is shorter than the ∼700 bp typically required for conventional non-CRISPR/Cas based approaches used for genetic modification in cyanobacteria (Behler et al., 2018). More importantly, for all target genes (*cax3-4*, *gor* and *sodB*), full chromosome segregation was achieved after the first patch. This is particularly significant as cyanobacteria are generally polyploid and several rounds of subculturing with increasing concentrations of antibiotics are required to achieve full segregation. CRISPR/Cas systems enhance the editing efficiency in cyanobacteria by promoting homology-directed repair, as most lack non-homologous end joining (Behler et al., 2018; Rajaram et al., 2020). Plasmid curing is essential for repeated genome editing in cyanobacteria due to limited antibiotic markers and slow growth, which makes multiple subculturing in antibiotic-free media time-consuming (Dias et al., 2015). Here, we hypothesize that selecting solely for the Sm^R^/Sp^R^ cassette encoded in the repair template and not for the Km^R^ marker on the editing plasmid during the initial selection step, combined with the associated toxicity of Cas12a, enriched for cells that had undergone both successful recombination and loss of the plasmid. Curing of Cas12a-based plasmid can be further accelerated though the use of *sacB* mediated sucrose counter-selection as described in (Niu et al., 2019).

Most, if not all, freshwaters cyanobacteria harbour a *sodB* gene that has been shown to be essential for cell growth of *Synechocystis,* which harbours no other SOD-encoding gene (Ke et al., 2014). Interestingly, we show that *sodB* is not essential for the growth of *Cyanothece* 7425 under standard phototrophic conditions. These results suggest that superoxide radicals can be handled by not only SodB, but also very likely by other scavengers yet to be identified. However, the cell death of *ΔsodB* mutant in presence of menadione clearly indicates that SodB plays an important role in the protection of *Cyanothece* 7425 against heightened oxidative stress. Similarly, the absence of Gor did not affect the growth of *Cyanothece* 7425, consistent with observations in both *E. coli* and yeast (Apontoweil and Berends, 1975; Muller, 1996). This suggests that either increased GSH synthesis or the thioredoxin reductase/thioredoxin system might compensate for the absence of GSSG reduction to GSH (Cameron and Pakrasi, 2010; Cassier-Chauvat et al., 2023; Prinz et al., 1997). The sensitivity of *Cyanothece* 7425 to cobalt, exacerbated in the *Δgor* mutant, is consistent with that observed in *Synechocystis* PCC 6803, where the glutathione system has been demonstrated to cope with cobalt-induced toxicity (Marceau et al., 2024; Morette et al., 2026). Indeed, in *Synechocystis* PCC 6803, knockout of GSH-dependent GST (Glutathione S-transferase) resulted in increased sensitivity to Co, accompanied by impaired Fe homeostasis and disruption of 4Fe-4S clusters (Marceau et al., 2024). Further investigation is necessary to better understand by which mechanisms cyanobacteria tolerate Co and the specific role of Gor in mitigating Co-induced toxicity.

The deletions to check whether *cax* genes are involved in iACC formation showed that at least Cax3 and Cax4 are not needed to make the inclusions. It is possible that the above-mentioned Cax1 and Cax2 proteins provide functional redundancy and compensate for the loss of both Cax3 and Cax4. Given the growth of iACC-forming cyanobacteria is highly Ca²⁺ dependent, Ca²⁺ trafficking is likely complex in these cyanobacteria (De Wever et al., 2019). Other transporters, such as Ca²⁺-associated P-type ATPases, can contribute to Ca²⁺ flux in *Cyanothece* 7425, and may modulate or mask the *Δcax3-4* mutant phenotype. Interestingly, a new gene family, *ccyA,* located in the same locus as *cax3-4* and adjacent to three putative P-type ATPases in *Cyanothece* 7425 (Fig. S2A), has been identified as a marker of iACC formation, and may play a role in Ca²⁺ transport and/or formation of CaCO_3_ inclusions (Benzerara et al., 2022). In the iACC-forming cyanobacterium *Microcystis aeruginosa* PCC 7806, genes encoding Cax also co-localized and co-expressed with *ccyA* (Bruley et al., 2025), suggesting their involvement in Ca²⁺ trafficking. Ongoing CRISPR/Cas12a mediated mutagenesis of these ATPases and the *ccyA* gene will help clarify their roles in iACC formation in *Cyanothece* 7425.

The CRISPR/Cas12a system has been demonstrated to be effective in generating gene knockouts and knock-ins, conditional mutants of essential genes, and large chromosomal deletions in model cyanobacteria (Niu et al., 2019; Ungerer and Pakrasi, 2016). Its utility has been successfully extended to two fast-growing cyanobacteria *Synechococcus* PCC 11801 (Sengupta et al., 2020) and *Synechococcus* PCC 11901 (Victoria et al., 2024). More recently, two non-model cyanobacteria, the unicellular nitrogen-fixing species *Cyanothece* ATCC 51142 and the filamentous *Leptolyngbya* sp. strain BL0902, have also found to be amenable to genetic manipulation using CRISPR/Cas12a to generate single/double gene knockouts (Bandyopadhyay et al., 2024; Gao et al., 2024). Consistent with these studies, we demonstrate here that the CRISPR/Cas12a system is an effective tool for genetic manipulation of the non-model cyanobacterium, *Cyanothece* 7425 endowed with several interests: intracellular biomineralization, metal toxicity tolerance, nitrogen fixation, etc. This study establishes a foundation for the future development of CRISPR/Cas12a-based strategies to generate other marker or marker-less mutants to harness the full potential of *Cyanothece* 7425.

## CONCLUSION

The biodiversity of non-model microorganisms plays a crucial role in expanding our understanding of fundamental metabolism and uncovering novel pathways and metabolites. Many of these organisms possess unique metabolic traits making them valuable resources for biotechnology and therapeutic applications. However, their effective study and exploitation depend on improving their cultivation under laboratory conditions and genetic tractability. In this context, we extended the use of CRISPR/Cas12a-based system to achieve targeted gene knockouts in the non-model cyanobacterium *Cyanothece* 7425. This bacterium is notable because of its ability to form iACC inclusions, a process relevant to the biogeochemical cycling of alkaline earth elements and to nuclear waste bioremediation. The establishment of these new genetic tools represents a significant step forward in elucidating the molecular mechanisms underlying iACC formation in cyanobacteria. Ongoing studies on other genes potentially involved in intracellular biomineralization will further our understanding of this fascinating process.

## Statements and Declarations

### Ethical Approval

Not applicable

### Author contributions

MAK: Conceptualization, Methodology, Investigation, Formal analysis, Writing- original draft & editing. AD, FSP, KB, CCC, FC: Formal analysis, Validation, Writing-review & editing. KB, CCC: Funding acquisition. SO: Conceptualization, Methodology, Investigation, Formal analysis, Supervision, Writing -review & editing

## Funding

This work was funded by the French National Agency (ANR): ANR Harley, ANR-19-CE44-0017, that also paid PhD fellowship of Monis Athar Khan.

### Availability of data and materials

All strains used in this study are available upon request from the corresponding authors.

## Acknowledgments

We are grateful to French National Agency for funding and to IMPMC microscopy platform at Sorbonne Université, France.

