## Supplementary Data for "Targeted genome editing of the non-model cyanobacterium *Cyanothece* PCC 7425 via CRISPR/Cas12a"

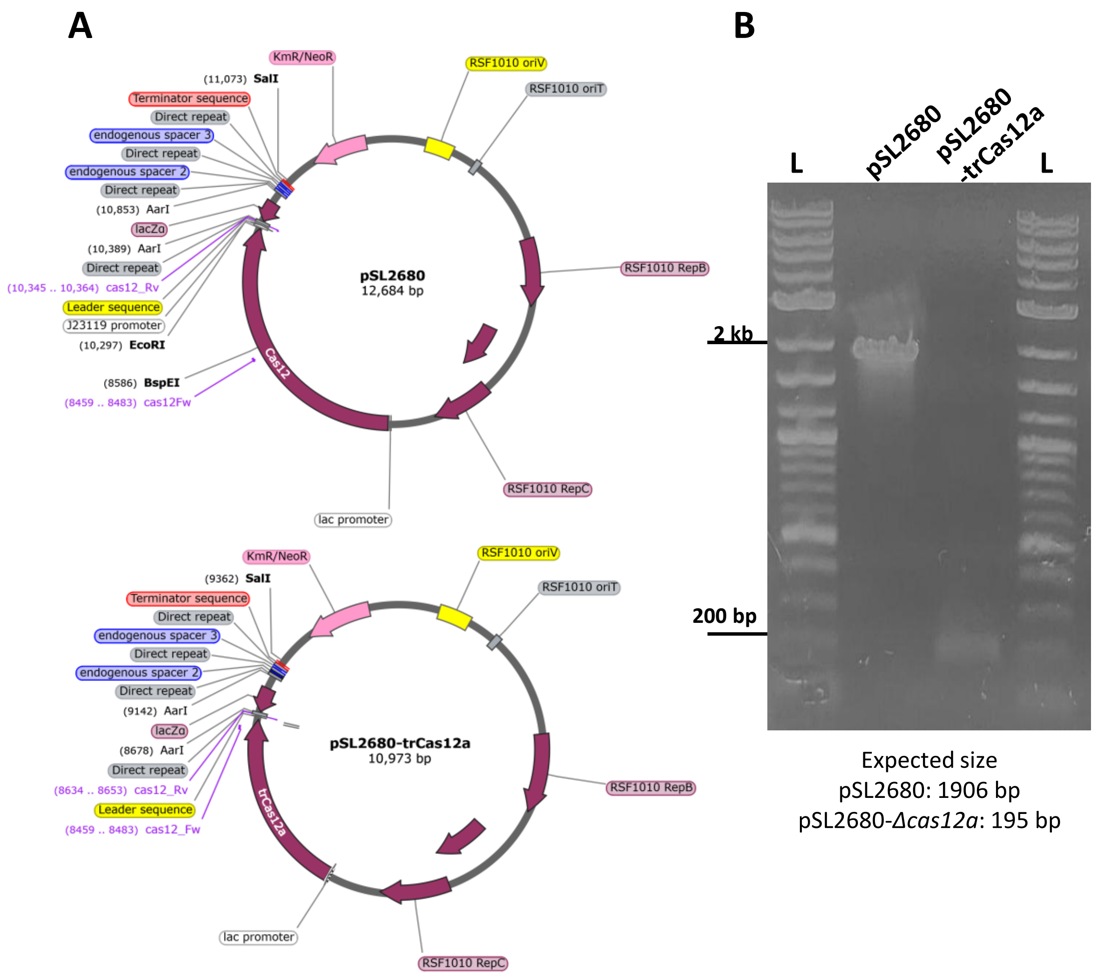


**Figure S1: (A)** The map of the pSL2680 (top panel) and pSL2680-trCas12a (bottom panel) plasmids, made using SnapGene. **(B)** PCR amplification of pSL2680 and pSL2680-trCas12a with Cas12-1 and Cas12-2 primers (Table S2) resolved on a 1% agarose gel indicating the corresponding product sizes. L denotes DNA ladder (1 kb plus, NEB).

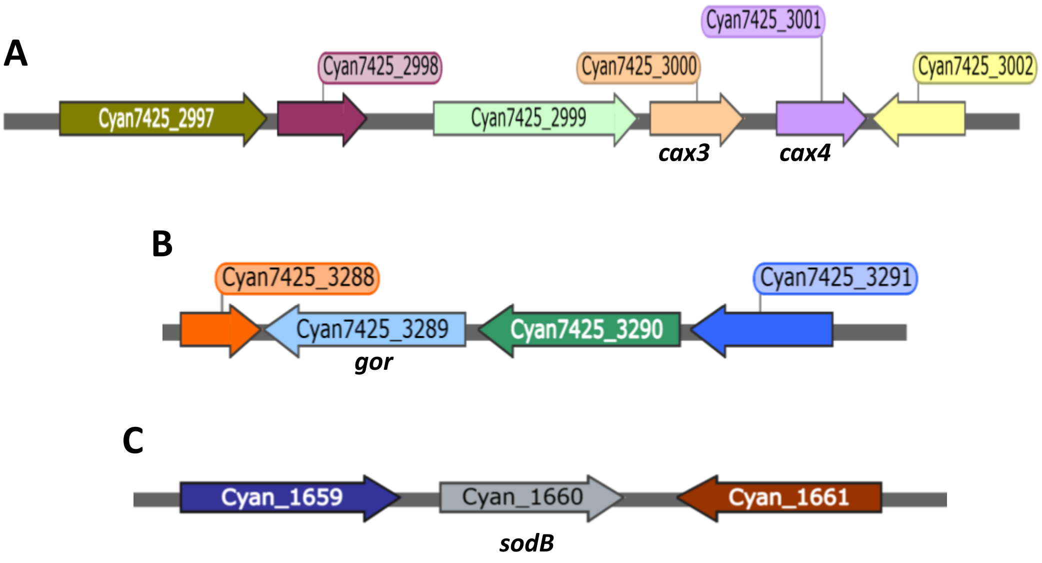


**Figure S2: (A)** Genomic region around the *cax3* (Cyan7425_3000) and *cax4* **(**Cyan7425_3001) genes. The upstream genes Cyan7425_2999 Cyan7425_2998 and Cyan7425_2997 encode a P-type ATPAse, protein of unknown function, and another P-type ATPAse, respectively. The downstream gene Cyan7425_3002 encodes an anion transporting ATPase. **(B)** Genomic region around the *gor* gene **(**Cyan7425_3289) showing its upstream gene Cyan7425_3288 (hypothetical protein and its downstream genes Cyan7425_3290 (nucleotide sugar dehydrogenase), Cyan7425_3291 (NAD-dependent epimerase). **(C)** Genomic region around the *sodB* gene (Cyan7425_1660) showing its flanking genes Cyan7425_1659 (hypothetical protein) and Cyan7425_1661 (dihydroorotase).

The genes are represented by large arrows pointing in their direction of transcription. The maps were made using SnapGene.

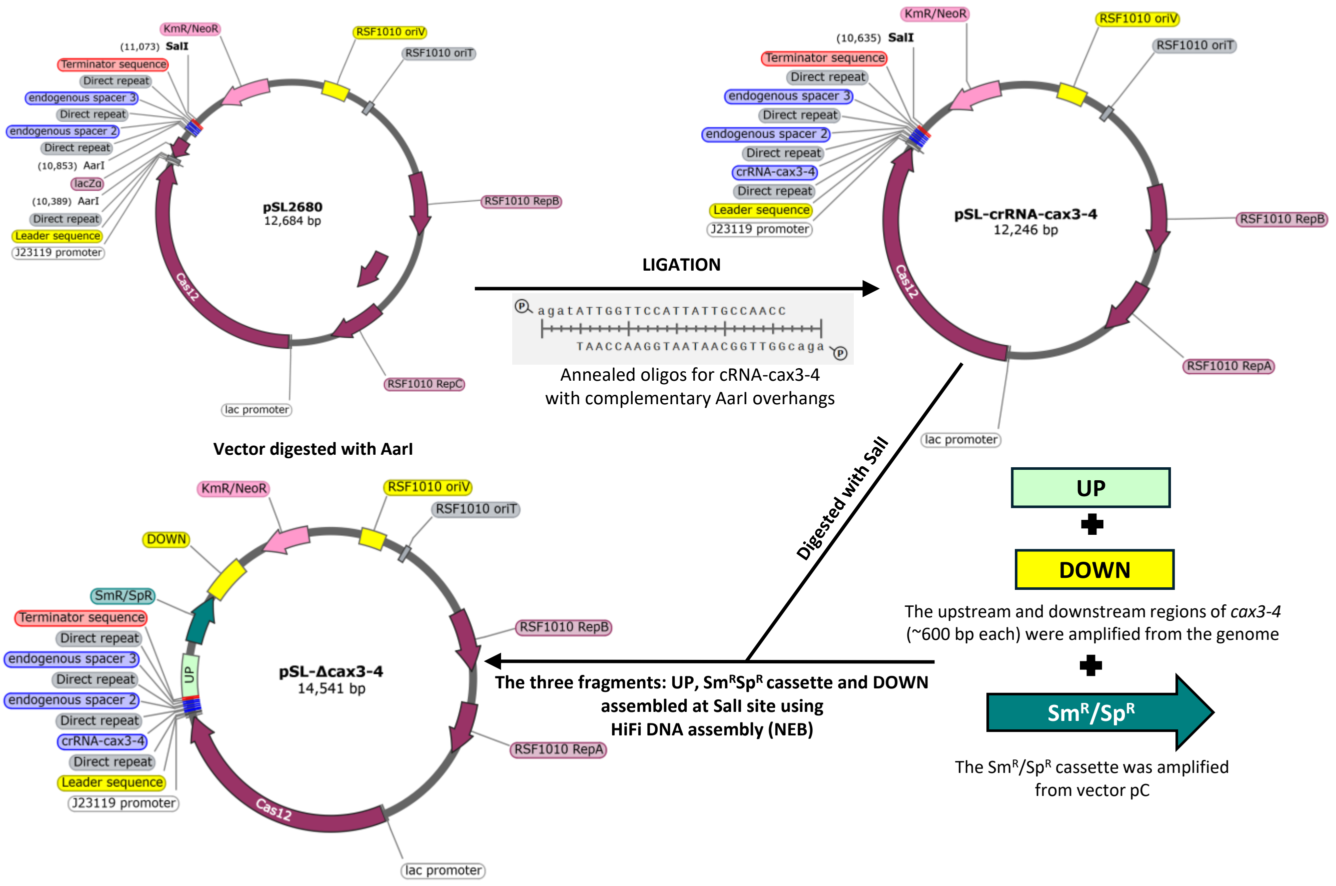


**Figure S3:** Schematic representation of the construction of pSL-*Δcax3-4* editing plasmid. The primers used are listed in Table S2. The pSL-*ΔsodB* editing plasmid was constructed similarly.

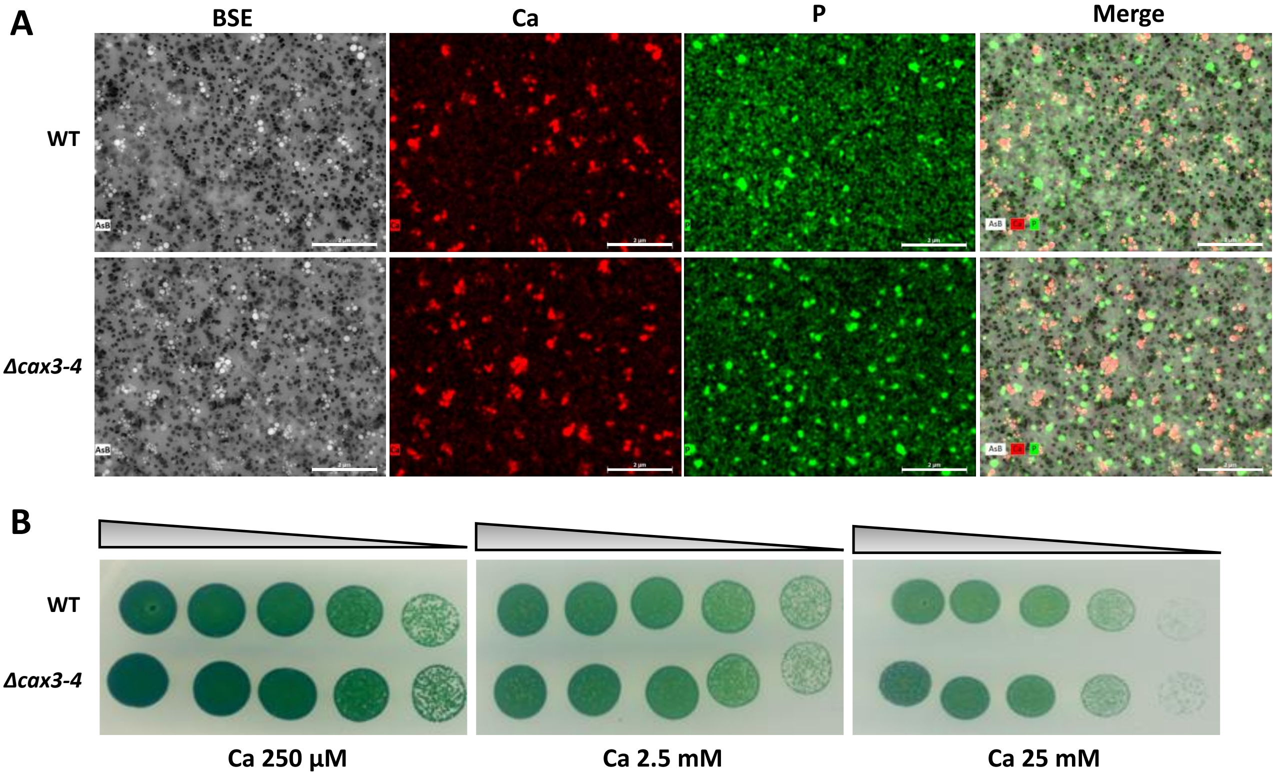


**Figure S4:** Phenotypic analysis of the *Δcax3-4* mutant. **(A)** Representative SEM images in BSE mode (leftmost panel) of WT and *Δcax3-4* cells grown in BG11 medium allowing detection of iACC inclusions (bright spots) and polyphosphate granules (light grey spots) as well as the pores (0.2 µm) of the filters (dark disks). The corresponding EDXS chemical maps (middle panels) of Ca and P are presented independently showing calcium-rich iACC inclusions as red dots and phosphate-rich PolyP granules as green dot, respectively. Merged images of the overlay of Ca and P chemical maps are shown in the rightmost panel. These representative images show no apparent difference in iACC inclusions between the WT and *Δcax3-4* mutant. Scale Bar: 2 µm **(B)** Spot-test assay for checking the influence of increasing calcium concentration on the growth of WT and *Δcax3-4* strains. Cultures adjusted to $OD_{750}$ = 0.1 were serially diluted 5-fold and spotted as 10 µL drops onto solid MM medium containing indicated calcium concentrations. The plates were incubated for 14 days at 30°C under standard light (2500 lux = 31.25 µmol photon $m^{-2}$ $s^{-1}$) prior to photography. No apparent differences were observed.

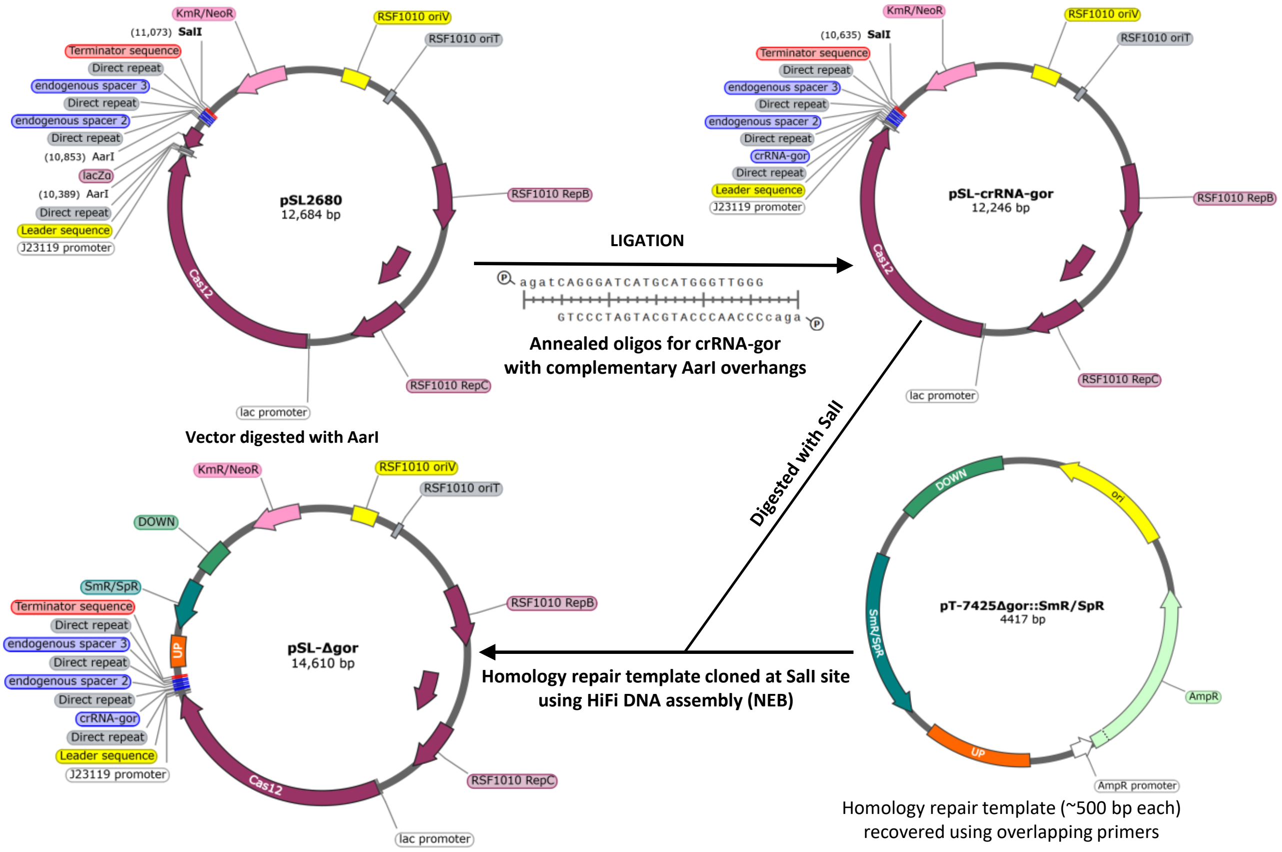


**Figure S5:** Schematic representation of the construction of pSL-*Δgor* editing vector. The primers used are listed in Table S2.

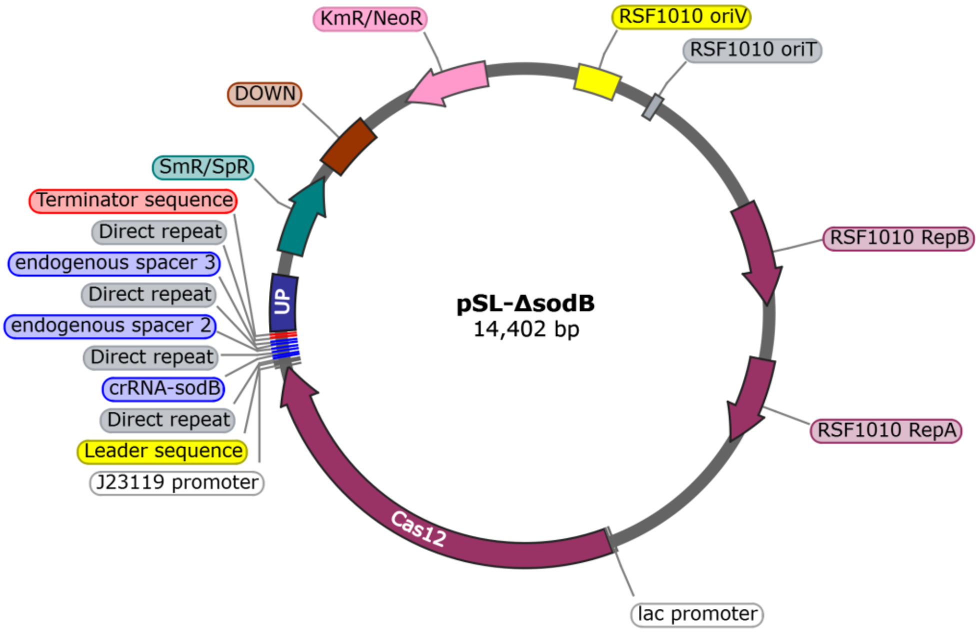


**Figure S6:** The map of pSL-*ΔsodB* editing vector made using SnapGene.

**Table S1:** List of plasmids used in this study.

| Plasmid | Features | Reference |
|---|---|---|
| pC | RSF1010 derivative harboring the λ phage promoter allowing strong constitutive expression. Used as donor of $Sm^R$/$Sp^R$ cassette | (Chenebault et al., 2020) |
| pSB2T | RSF1010 derivative harboring tac promoter. Used as control for conjugation experiments. | (Marraccini et al., 1993) |
| pSL2680 | RSF1010 derivative harboring the *cas12a* gene that is expressed from a lac promoter and the endogenous CRISPR array from *Francisella novicida* from a J23119 promoter. | (Ungerer and Pakrasi, 2016) |
| pSL2680-trCas12a | pSL2680 vector with truncated Cas12a (removal of amino acids 733-1330) that lacks the RuvC and nuclease domain | This study |
| pSL-*Δcax3-4* | pSL2680 vector with crRNA targeting *cax3* and $Sm^R$/$Sp^R$ cassette flanking homology platforms (~600 bp) for deletion of *cax3-4* genes | This study |
| pT-*Δgor* | pTwist plasmid with $Sm^R$/$Sp^R$ cassette flanking homology platforms for *gor* deletion | This study |
| pSL-*Δgor* | pSL2680 vector with crRNA targeting *gor* and $Sm^R$/$Sp^R$ cassette flanking homology platforms (~500 bp) for deletion of *gor* gene | This study |
| pSL-*ΔsodB* | pSL2680 vector with crRNA targeting *sodB* and $Sm^R$/$Sp^R$ cassette flanking homology platforms (~500 bp) for deletion of *sodB* gene | This study |

**Table S2:** List of primers used in this study.

| Primer Name | Sequence | Feature |
|---|---|---|
| gRNAseq-1 | TCTTGACAGCTAGCTCAGTCC | Verification and sequencing of pSL2680 plasmids |
| gRNAseq-2 | GACCGGCTACTAATACAAAAGG | |
| cas12_Fw | TTAGAATAAGAAATCATTCCACACA | Amplification of *cas12* gene on pSL2680 plasmid |
| cas12_Rv | CTTAGACCCGTTTTTGCCTA | |
| SmSp-1 | GATTTGCTGGTTACGGTGAC | Amplification of $Sm^R/Sp^R$ cassette |
| SmSp-2 | CCGACTACCTTGGTGATCTC | |
| **Construction of pSL-crRNA- Δcax3-4** | | |
| cr-cax3-4_Fw | agatATTGGTTCCATTATTGCCAACC | crRNA targeting *cax3* gene |
| cr-cax3-4_Rv | agacGGTTGGCAATAATGGAACCAAT | |
| F1-cax_Fw | AGCTTTAATGCGGTAGTTGGTACCGT CGACTGCCACCGACATTGCCCGA | ~600 bp upstream region from ATG of *cax3* gene |
| F1-cax_Rv | GCGTGAGCGCATAGATTGCCTCTGA ACGGTTCCG | |
| F2-cax_Fw | TCAGAGGCAATCTATGCGCTCACGC AACTGG | Recovery of $Sm^R/Sp^R$ cassette from pC |
| F2-cax_Rv | CCGGATTTGATCCGGCGTCGGCTTG AACG | |
| F3-cax_Fw | CAAGCCGACGCCGGATCAAATCCGG TCAGGATGGGGAAT | ~600 bp downstream region from stop codon of *cax4* gene |
| F3-cax_Rv | TGCCCGGATTACAGATCCTCTAGAG TCGACCTTCCCTGGGGGGACCGA | |
| **Verification of *Δcax3-4*** | | |
| cax-1 | ATTAATGACTCTGCCGCCCT | Amplification outside the upstream and downstream region of *cax3-cax4* gene |
| cax-2 | ATCTGGGCACGTTACGGTTA | |
| cax-3 | TTCTGTTGGGGATGCTAGCC | Amplification of *cax4* gene |
| cax-4 | CAGGGCAATTTGGAGACTGG | |
| **Construction of pSL-crRNA- Δgor** | | |
| cr-gor_Fw | agatCAGGGATCATGCATGGGTTGGG | crRNA targeting *gor* gene |
| cr-gor_Rv | agacCCCAACCCATGCATGATCCCTG | |
| F-pT-Δgor_Fw | AGCTTTAATGCGGTAGTTGGTACCGT CGAC CCAGTCACGACGTTGTAAAAC | Cloning of homology repair template from pT-Δgor to pSL- crRNA-Δgor |
| F-pT- Δgor_Rv | TGCCCGGATTACAGATCCTCTAGAG TCGAC TTCACACAGGAAACAGCTATGACCA T | |
| **Verification of *Δgor*** | | |
| gor-1 | ACAACTGGGTGTACCCTTGC | Amplification outside the upstream and downstream region of *gor* gene |
| gor-2 | ACAACTGGGTGTACCCTTGC | |
| gor-3 | TTCAGAAAGACCGACGGTAG | Amplification of *gor* gene |

| | | |
|---|---|---|
| gor-4 | GTCCTATGGAGCAAAAGTGG | |
| **Construction of pSL-crRNA- ΔsodB** | | |
| cr-sodB_Fw | agatAGTTTCACTATGGCAAGCACCA | crRNA targeting *sodB* gene |
| cr-sodB_Rv | agacTGGTGCTTGCCATAGTGAAACT | |
| F1-sodB_Fw | CGGTAGTTGGTACCGTCTCCGGTATGCCTGCCCT | ~500 bp upstream region from ATG of *sodB* gene |
| F1-sodB_Rv | AGCGCATAGGTGTTCTCCTTGTTTAACTGCT | |
| F2-sodB_Fw | AGAACACCTATGCGCTCACGCAACTGG | Recovery of $Sm^R/Sp^R$ cassette from pC |
| F2-sodB_Rv | CTAGCAGACCAGACTTGACCTGATAGTTTGGCTGTG | |
| F3-sodB_Fw | TCAAGTCTGGTCTGCTAGGCGTTACTACATCTCC | ~500 bp dowstream region from stop codon of *sodB* gene |
| F3-sodB_Rv | CGGATTACAGATCCTCTAGAGAAGGATGGGTTTAAAGATGCTTTTATCAAGC | |
| **Verification of *ΔsodB*** | | |
| sod-1 | CCGTAATTGTGCAGGTCGTC | Amplification outside the upstream and downstream region of *sodB* gene |
| sod-2 | CTGCAAATCAACGATCGCCT | |
| sod-3 | GCCTCCCTTACCCTACGATC | Amplification of *gor* gene |
| sod-4 | GGCAACAAAGTCCCAGTTCA | |
